# Harnessing genetic variations improving seedling vigor for successful crop establishment in deep sown direct-seeded rice

**DOI:** 10.1101/2023.03.28.534621

**Authors:** Nitika Sandhu, Ade Pooja Ankush, Jasneet Singh, Om Prakash Raigar, Sutej Bains, Taveena Jindal, Mohini Prabha Singh, Mehak Sethi, Gomsie Pruthi, Gaurav Augustine, Vikas Kumar Verma, Shivani Goyal, Aman Kumar, Harsh Panwar, Manvesh Kumar Sihag, Rupinder Kaur, Smita Kurup, Arvind Kumar

## Abstract

Improving seedling vigour remains key objective for breeders working with direct-seeded rice (DSR). To understand the genetic control of seedling vigour in deep sown DSR, combined genome-wide association mapping, linkage mapping, fine mapping, RNA-sequencing to identify candidate genes and validation of putative candidate genes was performed. Significant phenotypic variations were observed among genotypes in both F^3:4:5^ and BC^2^F^2:3^ populations. The mesocotyl length showed significant positive correlation with %germination, root and shoot length. The 3.79 Mb region on chromosome 7 reported to be associated with mesocotyl elongation. RNA-seq data and RT-PCR results identified and validated seven potential candidate genes. The potential candidate gene, *LOC_Os07g25150* codes for *Myb-30*-related transcription factor. The *LOC_Os07g25150* codes for *Myb 30*-related transcription factor, *LOC_Os07g17689* annotated as protein translation factor, *LOC_Os07g17770* as tryptophanyl-tRNA synthetase, *LOC_Os07g15440* as alanyl-tRNA synthetase family protein, *LOC_Os07g24100* as retrotransposon protein and the remaining others as expressed proteins. The antagonistic relationship between *Myb-30* and ethylene-mediated signalling (*EIN-3*) in regulating mesocotyl elongation have been observed. The functional characterization and knocking of *LOC_Os07g25150* codes for *Myb 30*-related transcription factor may provide better understanding of the mechanism behind mesocotyl elongation. The four promising introgression lines with longer mesocotyl length, longer root length and semi-dwarf plant height have been identified.

## Introduction

The mechanized direct-seeded rice cultivation is emerging as a natural resource-conserving, economically viable and climate smart strategy compared to the traditional puddled system of rice cultivation with huge potential to address the water-labour shortage and ensure sustainable rice production. The crop yield and resource-use efficiency mainly depend on the successful plant establishment in the field. The seed vigour defines the ability of the seeds to germinate and establish seedlings uniformly, rapidly, and robustly across variable environmental conditions. However, the poor emergence and non-uniform seedling establishment can lead to significant yield losses under DSR.

Improvement in the seedling vigour remains a breeding challenge in rice research (Finch-Savage et al., 2016) as it is not only important to enhance the crop yield but also can improve the crop resilience against the changing climatic conditions and the biotic impediments to rice yields. The sowing at recommended depth of 2 to 3 cm may lead to poor establishment and uneven crop stand due to heavy rain splashing, drought and heat stresses, more vapour pressure gradient, and predation (Kumar and Ladha, 2011; Yamauchi and Winn, 1996). In order to preserve the seeds and allow them to obtain moisture, deep sowing is an efficient strategy.

Deep seeding is generally not advised because it can be difficult for seedlings to penetrate the soil. Deep sowing can result in shallower crown placement, low dry mass build-up, late seedling emergence, and poor seedling establishment. One limiting factor is the elongation of mesocotyl and coleoptile, during seedling germination, the mesocotyl, an embryonic structure located between the coleoptile node and basal portion of the seminal root, is crucial in pushing the shoot tip across the soil surface. Particularly, high-yielding semidwarf rice varieties frequently have a short mesocotyl, which restricts their ability to emerge at deeper sowing depths. Thus, varieties with longer mesocotyl are useful to overcome issues faced by direct seeding. A longer mesocotyl will minimize sensitivity to seeding depth in drill seeding and improve seedling establishment. The environmental and genetic factors both played major role in mesocotyl and coleoptile elongation. The genetic studies showed that the mesocotyl length is under the control of multiple genes and could be stably inherited (Dilday et al., 1990).

The modern semidwarf cultivars have a short mesocotyl which does not allow crop establishment especially when seeds are drilled deeper in the soil and land levelling is not precise. The mechanistic knowledge of the basis of regulation of seed germination, seedling establishment and vigour from deeper sowing depth is limited. No doubt, the past domestication has provided an incremental improvement in seed characteristics but there is an urgent need to understand the regulatory mechanism behind the seed characteristics that serve as adaptive responses to the seed environment if sown at deeper soil depth. Many genetic loci associated with it have been reported (Regan et al., 1992; Redona and Mackill, 1996; Cui et al., 2022, Miura et al., 2001; Zhang et al., 2005; Fujino et al., 2008; Wang et al., 2010; Xie et al., 2014; Dang et al., 2014; Liu et al., 2014; Zhang et al., 2017; Zhao et al., 2019) but only a few have been validated and very few have been assessed for their impact mechanism on germination from depth. To breed rice varieties capable of germinating from depth under direct-seeding, there is need to explore the genomic regions or candidate genes for mesocotyl and coleoptile length that can be further used in molecular breeding by marker assisted selection.

The double digest restriction-site associated sequencing (ddRAD-seq) is a very flexible and cost-effective genotyping strategy providing in-depth insights into the genetic variations existing in the rice germplasm. A total of 39,137 polymorphic single nucleotide polymorphisms (SNPs) obtained through double digest restriction site associated DNA (dd-RAD) sequencing of 40 genotypes consisting of 20 rice landraces, and 20 released high yielding rice varieties used for association analyses revealed 188 stable SNPs associated with six traits across three environments. Mesocotyl and coleoptile length, along with seedling emergence and establishment are three important traits for determining high rice yields in DSR systems (Lee et al., 2017). It is critical to identify and exploit the genetic variations for seedling emergence from deep soil layer.

The next generation sequencing technologies such as dd-RAD provides SNP markers that can be utilized for the identification of alleles associated with the target traits (Poland and Rife, 2012) utilizing different mapping methods such as genome-wide association study (GWAS) and the QTL mapping. Recently, a number of studies used the combined GWAS and QTL mapping approaches to identify and validate the genomic regions/ marker-trait associations (MTAs) associated with complex traits (Sallam et al., 2022; Zhang et al., 2015; Gardiner et al., 2020). The GWAS is performed on a population constituting the unrelated individuals, while the QTL mapping used a biparental population (Alqudah et al., 2020; Zimmer et al., 2018).

Sufficient ground work in form of identification of donors, marker-trait association (MTAs)/QTL and segregating material as well as phenotyping platform tested on a wide scale is available. To understand the genetic control of rice seedling vigour under DSR, genome wide association studies for multiple seedling traits in 684 accessions from the 3000 Rice Genomes (3K-RG) population in both the laboratory and in the field at three planting depths (4, 8 and 10 cm) was carried out (Menard et al., 2021). Significant MTAs/QTL for mesocotyl and coleoptile length, percentage seedling emergence and shoot biomass in this panel were identified. The potential donors to be used in genomics-assisted breeding programs have been identified. The bi-parental mapping population using the identified donors from 3K-RGP subset (3,000 rice genome project) have been developed for the targeted traits. The present study designed to validate the identified genomic regions associated with seeding vigour from deep sowing depth under DSR. The study involves the genetic analysis of F_3_ and F_4_ mapping population derived from a cross between PR126 and IRGC 128442 for mesocotyl and coleoptile elongation/emergence from deeper soil depths and the development of near isogenic lines in background of PR126.

## Results

### Selection of plant material for the development of mapping population

A sub-set of 684 re-sequenced accessions including 266 tropical and temperate *japonica* 266 indica, 7 aromatic, 131 *aus/boro* and 14 admix subgroups from the 3K RGP representing the genetic and phenotypic diversity were selected. The phenotypic characterization of the accessions for seedling emergence traits at three planting depths (4, 8 and 10 cm) under field conditions and laboratory conditions was carried out. Genome wide association study (GWAS) showed a hotspot on the short arm chromosome 7 for mesocotyl and coleoptile length, percentage seedling emergence and shoot biomass in this panel (Menard *et al*., 2021). The seeds of the 6 selected accessions (Aus344, N22, Kula Karuppan, NCS237, Ashmber and IRGC 128442) were procured from IRRI and used as potential donors in genomics-assisted breeding program at PAU. The six accessions were evaluated first by drilling at 8 cm soil depth under dry direct seeded cultivation conditions under field conditions (Figure 1) and the best donor IRGC 128442 was chosen for the development of mapping populations in background of PR126.

**Figure 1.**
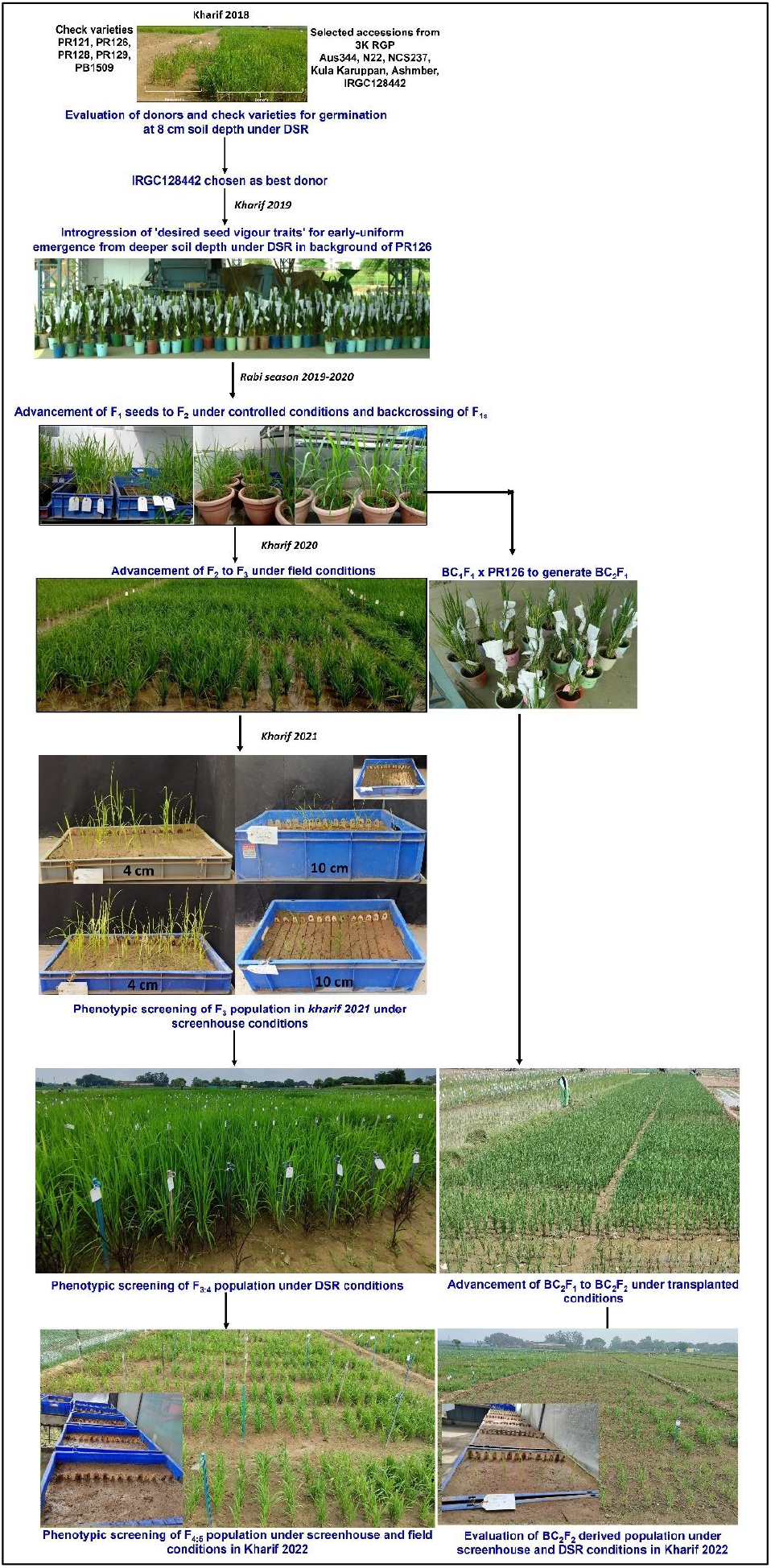
The details of the steps involved in development and evaluation of mapping populations across seasons under field and screenhouse conditions.

### Population development and phenotypic characterization

The complete steps involved in development and evaluation of mapping populations is presented in Figure 1. The four traits (%germination, mesocotyl and coleoptile length, root and shoot length) potentially linked to the improved germination and seedling vigor of rice from deep sowing depth and 5 traits {days to 50% flowering (DTF), plant height (PHT), panicle length (PL), number of panicles per plant (P/P) and grain yield (GY)} associated with grain yield were qualified in the present study. The significant phenotypic variations were observed among genotypes across different depths under screenhouse and field conditions in both F_3:4:5_ and BC_2_F_2:3_ populations (Table 1). The frequency distribution of the traits measured in the present study were determined for each year at different sowing depths under both screenhouse and field conditions. Most of the traits segregated continuously and almost fitted a normal distribution under different depths, years and conditions (Supplementary Figures S1, S2). Compared to 10 cm sowing depth, better germination was observed under 4 cm sowing depth.

**Table 1.**
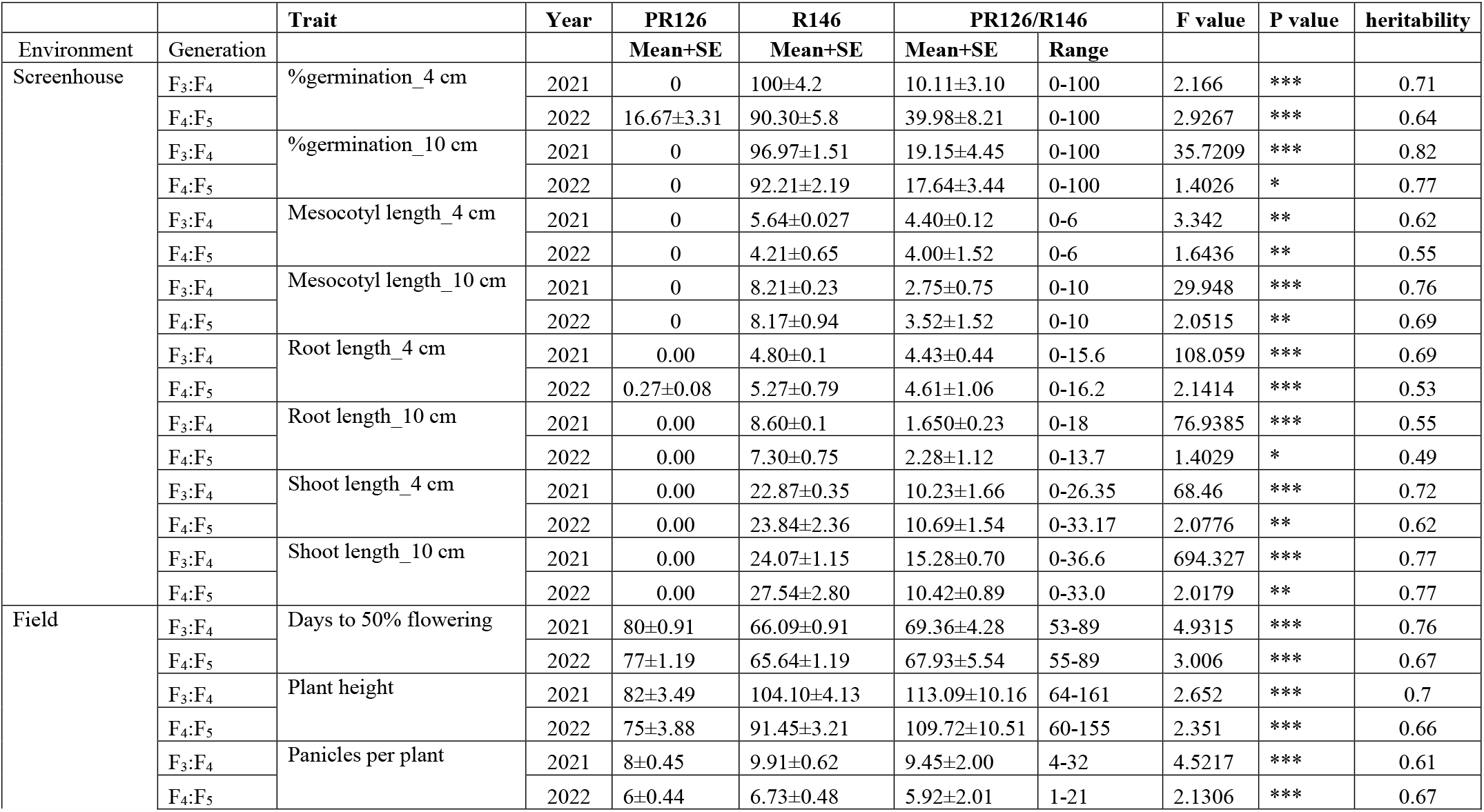
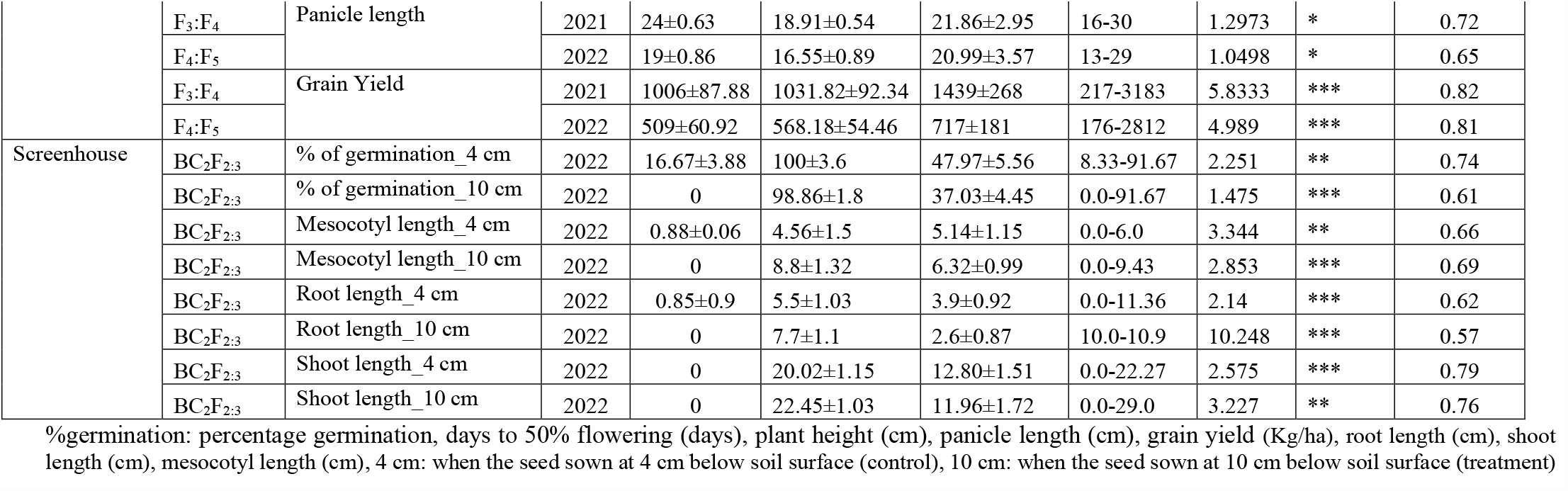
Detailed information on analysis of variance (ANOVA), mean, range and heritability of the traits measured across years in different growing environments in both F_3:4:5_ and BC_2_F_2:3_ mapping populations in background of PR126.

The % germination and mesocotyl and coleoptile length of IRGC 128442 at 4 cm and 10 cm sowing depth were significantly higher than the recipient parent (PR126). IRGC 128442 showed early flowering compared to PR126. The plant height of IRGC 128442 was significantly higher than the PR126 across years. The mean and range values of the traits measured in both F_3:4:5_ and BC_2_F_2:3_ populations under screenhouse and DSR field conditions in 2021 and 2022 are presented in Table 1. The correlation between different traits at different depths considering the pooled mean data is presented in Figure 2a. The mesocotyl length at 4 cm sowing depth found to be significantly and positively correlated with root length (r = 0.85, p <0.001) and shoot length at 4 cm (r = 0.93, p <0.001) (Figure 2a). Similarly, mesocotyl length at 10 cm sowing depth showed significantly positive correlation with root length (r = 0.91, p <0.001) and shoot length at 10 cm (r = 0.96, p <0.001) (Figure 2a). The phenotypic correlation coefficient analysis showed that the %germination at 4 and 10 cm of sowing depth is significantly and positively correlated with mesocotyl length at 4 (r = 0.80, p <0.001) and 10 cm (r = 0.69, p <0.001) of sowing depth, respectively (Figure 2a).

**Figure 2.**
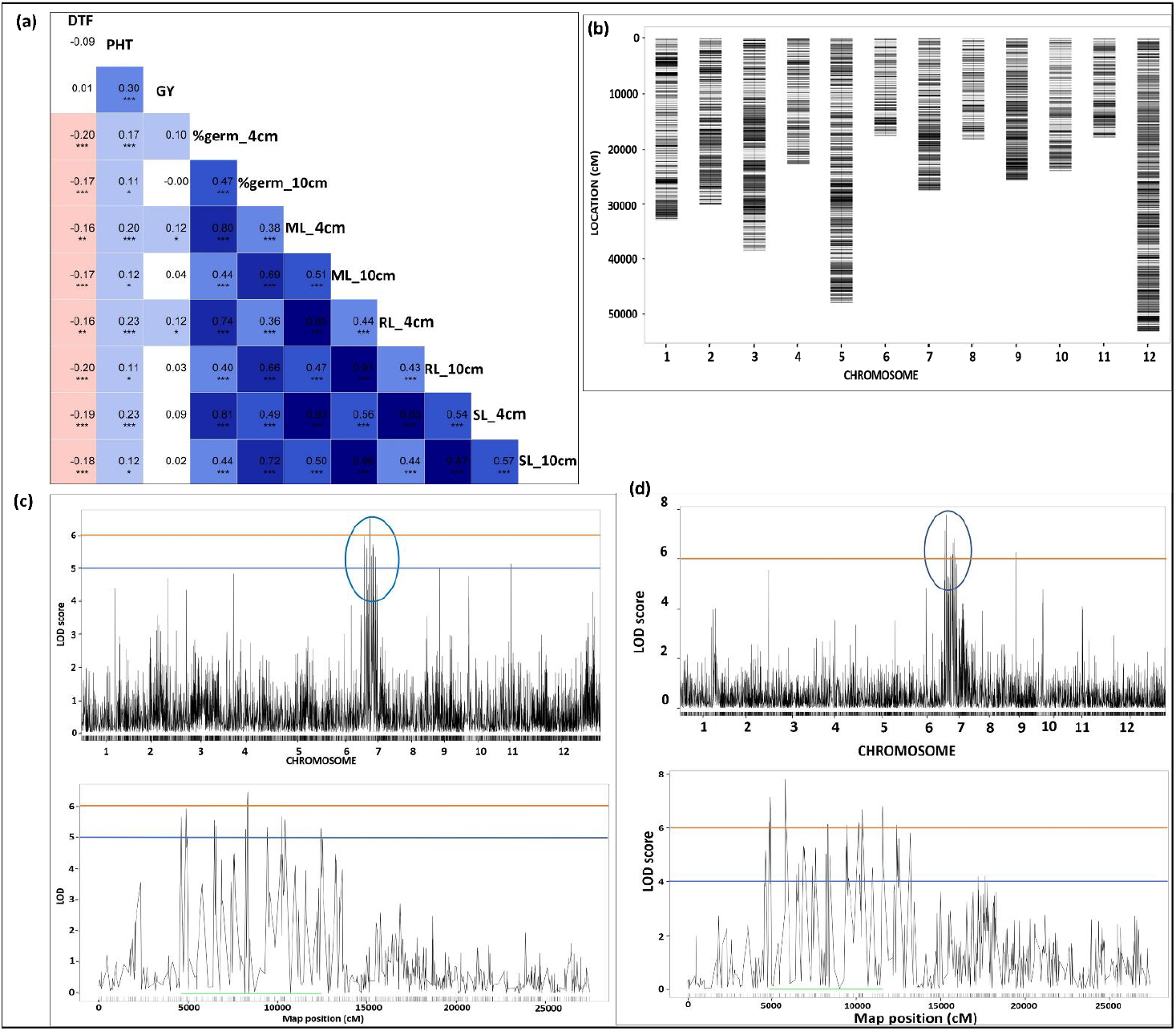
**(a)** Plots of Pearson’s *r-values* showing the correlation among traits measured at 4 and 10 cm of sowing depths. The blue colour indicates positive correlation and red colour indicated the negative correlation among different traits, the variation in colour intensity is representing the strength of the correlation among the traits. *Significance at <5% level, **significance at <1% level, ***significance at <0.1% level (**b)** SNP density plot showing the distribution of SNPs along the chromosomes **(c)** QTL likelihood curves of LOD scores for %germination at 10 cm sowing depth considering pooled mean analysis across years on chr 7 in PR126 x IRGC 128442 mapping population **(d)** QTL likelihood curves of LOD scores for mesocotyl length at 10 cm sowing depth considering pooled mean analysis across years on chr 7 in PR126 x IRGC 128442 mapping population

### SNP filtering and Linkage map construction

A total of 5,256,378 variants [SNPs+Indels], 4,824,612 SNPs and 4,779,408 biallelic SNPs were identified in the PR126/IRGC 128442 mapping population and in the parental lines. The filtering was carried out considering parameters such as minor allele frequency (MAF), genotype depth, breeding lines and allele missingness and parental polymorphism. After filtering, the number of SNPs reduced to 11,196. Of the 420 breeding lines, 350 F_3:4:5_ breeding lines were retained after filtering the breeding lines with high genotype missingness. Next, removing 898 SNPs that showed segregation distortion (p < 0.01), finally generated a set of 10,298 SNPs. The SNPs were finally grouped on 12 chromosomes (Figure 2b). The genetic linkage map spanned 1561.609 cM with an average SNP interval of 0.2075 cM (Table S1). The genetic length of each linkage group (LG) ranged from 97.91 (LG11) to 172.2 cM (LG1), with an average SNP distance of 0.21 to 0.19 cM (Supplementary Table S1).

### QTL mapping and fine mapping

To identify the genomic regions associated with traits improving germination and seedling vigor and grain yield/yield associated traits under DSR, QTL mapping study was performed on a set of 350 F_3:4:5_ breeding lines using 10,298 SNPs. In our previous genome wide association mapping, we found that the main QTL for mesocotyl and coleoptile length, percentage seedling emergence and shoot biomass in the 684 accessions selected from 3K-RGP (rice genome project) are co-located on 4.84 Mb region on the short arm of chromosome 7 (Menard *et al*., 2021). The earlier identified genomic regions in the 3K-RG subset using genome wide association mapping approach were validated in the present study on F_3:4:5_ mapping population developed using the donor (IRGC 128442) selected from 3K-RG subset in background of popular *parmal* rice variety (PR126) grown in Punjab. The genomic region extended from 8.83 Mb to 15.44 Mb (6.61 Mb) on chromosome 7 showed association with % germination, mesocotyl length (Table 2, Figs. 2c, 2d), root and shoot length at different sowing depths across seasons (Table 2, Supplementary Figure S3). The striking finding of the present study is that there is a convergence of genomic region associated with %germination and mesocotyl and coleoptile length on chromosome 7 using QTL mapping approach with the genomic region identified earlier on chromosome 7 using genome wide association mapping approach (Figure 3). Concerning, the grain yield and yield associated traits, a genomic region spanning 8.99 Mb region on chromosome 1 reported to be associated with DTF, PHT and GY traits (Table 2, Supplementary Figure S4).

**Table 2.** Detailed information on the genomic regions associated with different traits of interest identified in the different mapping populations.

| Mapping population | Trait | Chr | QTL interval (bp) | QTL span (Mb) | Left flanking marker | Peak marker (bp) | Right flanking marker | LOD | R<sup>2</sup> |
| --- | --- | --- | --- | --- | --- | --- | --- | --- | --- |
| **F<sub>3:4:5</sub>** | %germination_4 cm | 7 | 8906211-14769869 | 5.86 | 7:8906211 | 7:10322112 | 7:14769869 | 10.10 | 12.43 |
|  | %germination_10 cm | 7 | 8831295-14769869 | 5.94 | 7:8831295 | 7:12711103 | 7:14769869 | 5.60 | 7.10 |
|  | Mesocotyl length_4 cm | 7 | 8906211-15425220 | 6.52 | 7:8906211 | 7:11758772 | 7:15425220 | 9.90 | 12.19 |
|  | Mesocotyl length_10cm | 7 | 8831295-15438381 | 6.61 | 7:8831295 | 7:9530368 | 7:15438381 | 7.70 | 9.58 |
|  | %germination_4cm | 2 | 25640406-30681013 | 5.04 | 2:25640406 | 2:26193032 | 2:30681013 | 7.80 | 9.77 |
|  | %germination_4cm | 9 | 8088106-8164448 | 0.08 | 9:8088106 |  | 9:8164448 | 6.30 | 7.93 |
|  | Mesocotyl length_4 cm | 10 | 4770677-4770755 | - | 10:4770677 | 10:4770686 | 10:4770755 | 6.10 | 7.66 |
|  | Mesocotyl length_4 cm | 11 | 15550458-15550470 | - | 11:15550458 | 11:15550468 | 11:15550470 | 4.70 | 5.95 |
|  | Root length_4cm, Shoot length_4cm | 7 | 8906211-10369837 | 1.46 | 7:8906211 | 7:10369837 | 7:10369837 | 5.20 | 11.93 |
|  | Root length_10cm | 3 | 547685-1730359 | 1.18 | 3:547685 | 3:777138 | 3:1730359 | 8.80 | 10.88 |
|  | Days to 50% flowering, Plant height, Grain yield | 1 | 25344969-34329979 | 8.99 | 1:25344969 | 1:28134927 | 1:34329979 | 11.80 | 14.41 |
|  | Days to 50% flowering, | 3 | 858902-1730359 | 0.87 | 3:858902 | 3:894084 | 3:1730359 | 9.30 | 11.51 |
|  | Days to 50% flowering, | 8 | 6986267-7377701 | 0.39 | 8:6986267 | 8:7232778 | 8:7377701 | 7.20 | 9.03 |
|  | Days to 50% flowering, | 9 | 8088106-8088106 | - | - | 9:8088106 | - | 6.20 | 7.82 |
| **BC<sub>2</sub>F<sub>2:3</sub>** | %germination_4 cm | 7 | 10923664 | - | - | 10923664 | - | 3.30 | 9.6 |
|  | %germination_10 cm | 7 | 10923664 | - | - | 10923664 | - | 3.00 | 10.2 |
|  | Mesocotyl length_4 cm | 7 | 10923664-14713452 | 3.79 | K_10923664 | K_14713452 | K_14713452 | 8.68 | 11.28 |
|  | Mesocotyl length_10cm | 7 | 10923664-14713452 | 3.79 | K_10923664 | K_14713452 | K_14713452 | 9.00 | 12.80 |
|  | Shoot length_10 cm | 7 | 10923664-14713452 | 3.79 | K_10923664 | K_14713452 | K_14713452 | 3.30 | 11.20 |

**Figure 3.**
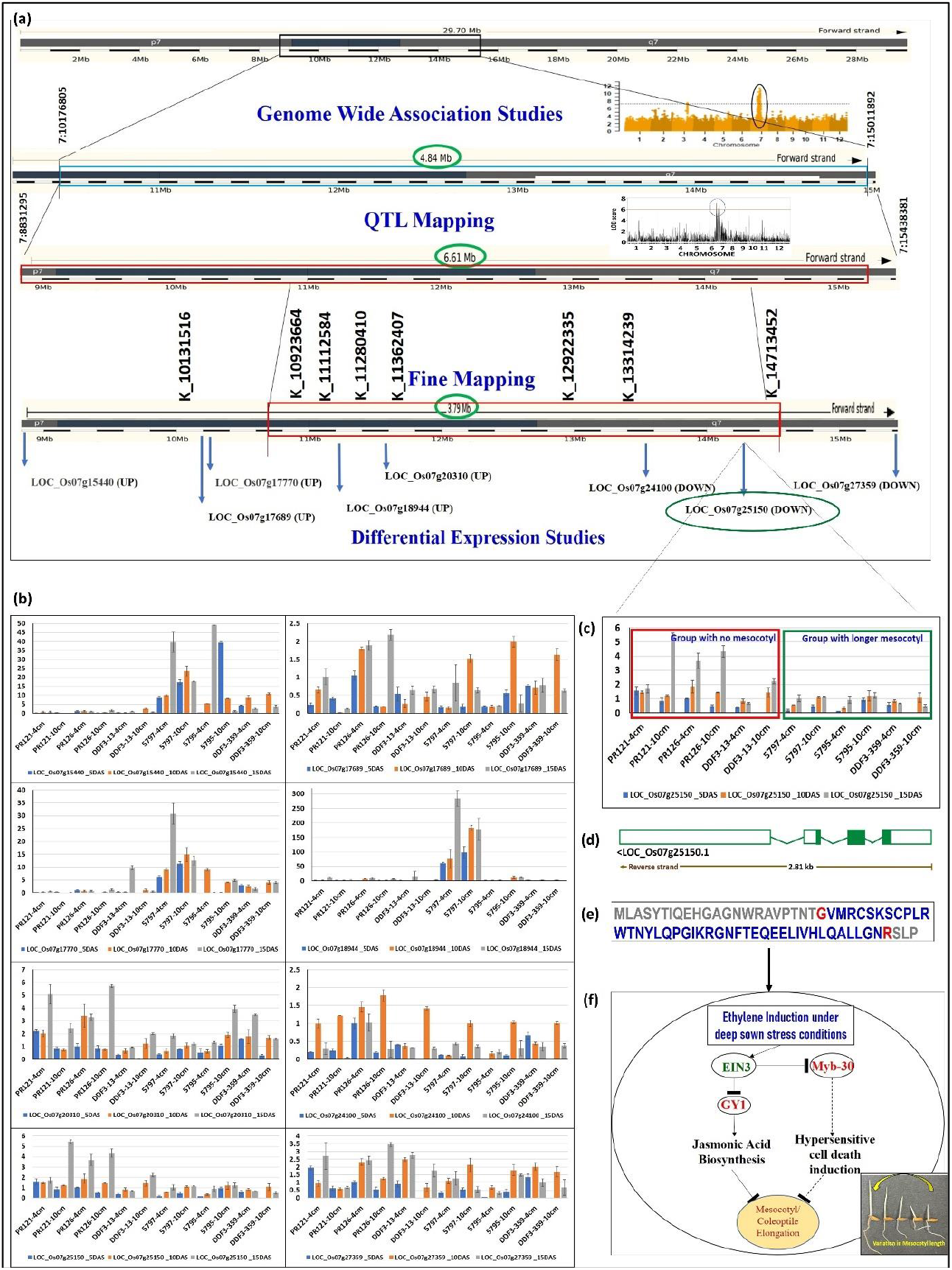
**(a)** The combined approaches involving genome-wide association mapping, linkage mapping and fine mapping followed by transcriptome based RNA-seq analysis identifying genomic region and putative candidate genes associated with mesocotyl elongation on chromosome 7 under deep sown direct-seeded rice cultivation conditions (**b)** validation of 8 putative candidate genes by RT-qPCR studies **(c)** validation of *LOC_Os07g25150* on a panel of 6 genotypes constituting 3 with longer and 3 with short/no mesocotyl length by RT-qPCR studies **(d)** gene structure of the validated candidate gene *LOC_Os07g25150* (codes for Myb-30 transcription factor) **(e)** protein sequence of the validated candidate gene LOC_Os07g25150 (codes for Myb-30 transcription factor) **(f)** Interplay between Ethylene induced *EIN-3* and Myb30 regulates mesocotyl elongation.

The backcross generation was finally applied to fine map the genomic region associated with traits improving germination of rice under DSR. The 15 KASP markers which were designed in the 6.6 Mb genomic region on chromosome 7, were first checked for parental polymorphism. A total of 8 KASP assays showed satisfactory results on parents used to genotype the BC_2_F_2:3_ plants (Figure 4a). Based on the genotypic analysis and phenotypic analysis of 339 BC_2_F_2:3_ plants, the genomic region associated with %germination showed association at peak marker located at 10.92 Mb region on chromosome 7 and with mesocotyl and coleoptile length to 10923664 to 14713452 Mb (3.79 Mb) region (Figure 3).

**Figure 4.**
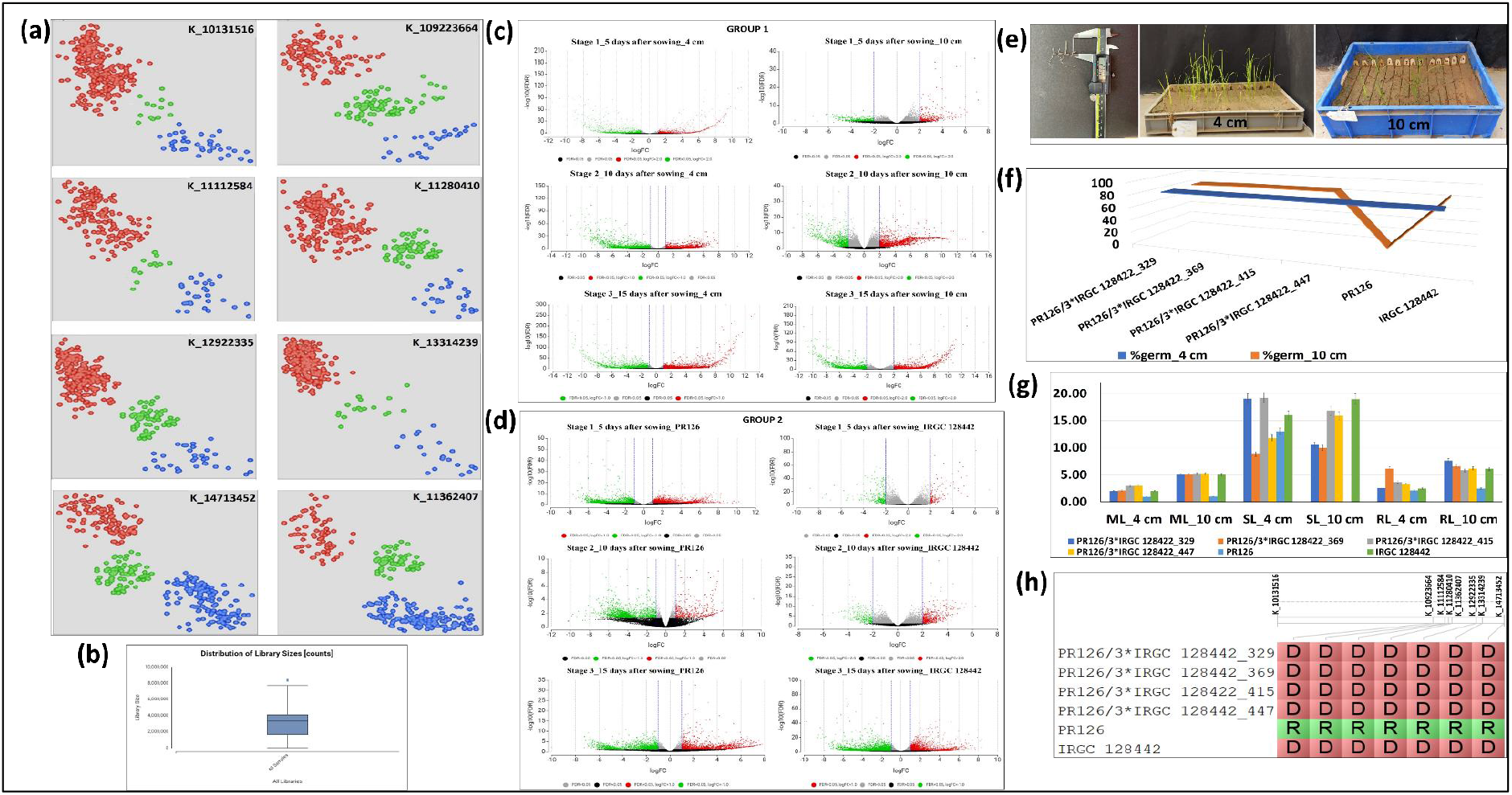
**(a)** The pictorial representation of the 8 KASP assays conducted on the 339 BC_2_F_2:3_ plants including parents. Blue color indicates the donor allele, red color indicates the recipient allele and green color indicates the heterozygotes. **(b)** The schematic representation of the library size for 106 samples used to generate RNA-seq data **(c)** The volcano plot showing differentially expressed genes in group I **(d)** The volcano plot showing differentially expressed genes in group II **(e)** The pictorial representation of the measurement of mesocotyl length using vernier calliper, phenotypic screening of the backcross population under control (4 cm) and deep sown (10 cm) conditions **(f)** The performance of selected promising breeding in comparison to recipient parent (PR126) and donor parent (IRGC 128442) lines in terms of mesocotyl elongation (ML), root length (RL) and shoot length (SL) under control (4 cm) and deep sown (10 cm) conditions **(g)** The performance of selected promising breeding in comparison to recipient parent (PR126) and donor parent (IRGC 128442) lines in terms of %germination under control (4 cm) and deep sown (10 cm) conditions **(h)** The allelic constitution of the selected breeding lines, recipient parent (PR126) and donor parent (IRGC 128442) in the fine mapped region on chromosome 7. D: donor parent allele, R: recipient parent allele

### Application of RNA-Sequencing to filter DEGs

To identify the differentially expressed genes between IRGC 128442 and the control variety PR126, a total of 108 RNA sequencing libraries were developed for both the PR126 and IRGC 128442 sown at 2 different depths (4 cm and 10 cm), and 3-time points (5 DAS, 10 DAS, and 15 DAS) with each having 3 biological and 3 technical replicates. The library size for 106 samples was distributed from 1665409 to 7727934 with a median value of 4090419 (Figure 4b). Two samples with a library size of 0 were discarded and were not further considered for the downstream analysis. Gene-level quantification analysis based on alignment with the reference genome annotation file was done for all the samples to generate a count table for differential expression analysis. The alignment results showed that an average of 93.45% reads for all 106 samples uniquely aligned to the *Oryza sativa* reference genome (Supplementary Table S2).

To compare the samples of the same rice cultivar under different treatment conditions [control (4cm) and stress (10cm)] and different rice cultivars (PR126 and IRGC 128442) under the same treatment at 3 different time points; a total of 2 groups were created with each having 6 subgroups. In group I, differential expression analysis between (IRGC 128442 vs PR126) was done at (Stage 1_5DAS - 10cm, Stage 2_10DAS - 10cm, Stage 3_15DAS - 10cm, Stage 1_5DAS - 4cm, Stage 2_10DAS - 4cm, Stage 3_15DAS - 4cm) subgroups, and in group II, differential expression among 2 treatments [control (4cm) and stress (10cm)] for the same rice cultivar was observed at (Stage 1_5DAS – IRGC 128442, Stage 2_10DAS – IRGC 128442, Stage 3_15DAS – IRGC 128442, Stage 1_5DAS - PR126, Stage 2_10DAS - PR126, Stage 3_15DAS - PR126) subgroups. The differentially expressed genes (DEGs) identified by RNA sequencing in both groups are summarized as a volcano diagram by estimating the expression of genes (Figures 4c, 4d). Eight differentially expressed genes; *LOC_Os07g15440, LOC_Os07g17689, LOC_Os07g17770, LOC_Os07g18944, LOC_Os07g20310, LOC_Os07g24100, LOC_Os07g25150* and *LOC_Os07g27359* were identified in the above QTL regions from both groups (Figure 3a). The expression (FPKM value) of 8 genes in 106 samples was analyzed to check whether the gene expression is consistent in three biological replicates and their corresponding 3 technical replicates. Out of the eight genes, 5 genes were upregulated at both Stage 2 and Stage 3 (*LOC_Os07g15440, LOC_Os07g17689, LOC_Os07g17770, LOC_Os07g18944* and *LOC_Os07g20310*), 1 gene was downregulated at both stage 1 and stage 2 (*LOC_Os07g27359*), 2 genes were downregulated at stage 3 (*LOC_Os07g24100*, and *LOC_Os07g25150*).

### Validation of the selected differentially expressed genes by RT-qPCR

To confirm the accuracy and reproducibility of transcriptome results of the above eight DEGs, qRT-PCR was performed using a panel of 6 samples (PR126, PR121, a line with very short mesocotyl from the PR126/IRGC 128442 mapping population, IRGC 128442, NCS237, a line with long short mesocotyl from the PR126/IRGC 128442 mapping population) (Figs. 3b, 3c). The results showed that the expression of seven genes was significantly different in the contrasting genotypes. The qRT-PCR analysis results revealed the relative trends in the expression patterns of the seven genes that were consistent with the RNA-sequencing data.

### Functional annotation of the validated genes

Out of the seven validated candidate genes, the potential candidate gene, the potential candidate *LOC_Os07g25150* gene transcript is 2329 bp in size, constitute four exons (Figure 3d) and the protein sequence of 68 amino acids (Figure 3e). The *LOC_Os07g25150* codes for *Myb 30*-related transcription factor (Figure 3f), *LOC_Os07g17689* annotated as protein translation factor, *LOC_Os07g17770* as tryptophanyl-tRNA synthetase, *LOC_Os07g15440* as alanyl-tRNA synthetase family protein, *LOC_Os07g24100* as retrotransposon protein and the remaining others as expressed proteins.

### Selection of promising lines

The four promising introgression lines from BC_2_F_2:3_ populations with improved %germination (Figures 4e, 4f), longer mesocotyl length, long root length, semi-dwarf seedling height (Figure 4g) and possessing the favourable allele combinations (Figure 4h) have been selected for further evaluation.

## Discussion

Keeping water-food security as well as farmer’s livelihood in mind, technologies aimed at saving water-labor-energy such as DSR system hold significance. However, poor emergence and inadequate seedling establishment led to substantial yield loss under DSR. The rice breeding programme at PAU has taken initiative to identify and characterize potential donors of improved seedling vigor and started backcross breeding to transfer and consolidate these traits with the help of precise phenotypic screening as well as use of molecular markers linked to these traits.

Due to the high resolution and dense genome coverage, many genomic regions associated with complex polygenic traits can be detected in GWAS compared to the biparental QTL mapping (Sallam et al., 2022). However, the core weakness of the GWAS mapping is the high false-positive rates resulted from the population structure (Larsson et al., 2013). The consideration of the principal component analysis (PCA), the structure (Q) as a fixed effects and the kinship matrix (K) (Mahuku et al., 2016) may solve the problem resulted in the rare to few false positives in the GWAS studies. In contrast, the QTL mapping has not the problem of the presence of the population structure. To understand the genetic control of rice seedling vigour under DSR; combined GWAS, QTL mapping and RNA-sequencing to identify candidate genes and KASP markers and validation of putative candidate genes have been performed in the present study. The integration of different genomic technologies is a perfect way to dissect the genetic basis of complex traits.

The GWAS for multiple seedling traits in 684 accessions from the 3000 Rice Genomes (3K-RG) was carried out by Menard et al., (2021). The significant MTAs/QTL for mesocotyl length, coleoptile length, seedling emergence and shoot biomass were identified in 4.84 Mb on chr 7 and validated on F_3:4:5_ mapping population developed using donor (IRGC 128442, selected from 3K-RG) in background of popular *parmal* rice variety (PR126) using linkage mapping 8.83-15.44 Mb (6.61 Mb)}. The backcross generation was applied for fine mapping {10923664 to 14713452 Mb (3.79 Mb)}. The eight candidate genes were selected from the RNA-seq data and seven genes were validated in the RT-PCR expression studies. The 8 putative candidate genes reported to be associated with cellular response to DNA damage, hypersensitive cell death response, ubiquitination/deubiuitination, aminoacylation, metal-ion binding, membrane, nucleus, cytoplasm, and mitochondria (Supplementary Figure S5).

Out of the 7 validated potential candidate genes in response to deep sowing for mesocotyl elongation in the main QTL interval, the 4 genes (*LOC_Os07g18944, LOC_Os07g20310, LOC_Os07g24100* and *LOC_Os07g25150*) were found in the fine mapped genomic region and 3 genes in upstream region *(LOC_Os07g15440, LOC_Os07g17770, LOC_Os07g17689*). The *LOC_Os07g25150* codes for a *Myb 30*-related transcription factor protein which binds specifically to 5’-AACAAAC-3’ DNA sequence (Li et al., 2009) and has been validated for initiating the hypersensitive cell death response in Arabidopsis (Daniel et al., 1999). Ethylene insensitive-3 (EIN-3) is a major factor regulating mesocotyl elongation by downregulating the *GY1* gene and inhibiting the jasmonic acid pathway (Xiong et al., 2017) (Figure 3f). EIN3 has been found to directly interact with Myb 30 transcription factor and hider its downstream cascade (Xiao et al., 2019) (Figure 3f). This antagonistic relationship between Myb-30 and EIN-3 in regulating mesocotyl elongation still remains unexplored. Our RNA-seq data and RT-PCR results suggest similar trends between *EIN-3* and *Myb-30* in the genotype possessing longer mesocotyl. This implies a strategic role and cross-talk between ethylene-mediated signalling pathway along with *Myb-30* in regulating mesocotyl elongation.

The knockout of *Myb-30* gene may help to better understand the mechanism behind mesocotyl elongation. The knocking of candidate gene may provide a powerful way to mesocotyl elongation improving seedling emergence, seedling vigor and providing adaptation to rice under DSR. The knockout mutation in the negative regulator of gene may lead to the development of DSR adapted rice varieties with longer mesocotyl length as well as possessing the capability to germinate from deeper soil depth. As the best of our knowledge, four genes for mesocotyl elongation, OsGY1 (Xiong et al., 2017), OsGSK2 (Sun et al., 2018), OsSMAX1 (Zheng et al., 2020), and OsPAO5 (Lv et al., 2021) have been cloned. None of the genes located on same position as the genomic region we have detected on chr 7. The seedling vigour trait reported to be associated with plant height, increase in plant height may lead to lodging under DSR. The introgression lines possessing alleles associated with longer mesocotyl length and semi-dwarf plant type have been identified. The identified promising lines may provide improvements in seedling establishment over currently available un-adapted rice varieties with water-labour saving. Farmers does not need to increase seed rate or spend more money to compensate poor seedling establishment under DSR.

## Conclusion and future perspectives

The integrated approach involving GWAS, QTL mapping and fine mapping led to the identification of 3.79 Mb region on chromosome 7 to be associated with mesocotyl elongation. A total of 7 potential genes including 4 genes in the fine mapped region and 3 genes in the upstream stream have been identified. The potential candidate gene, *LOC_Os07g25150* codes for *Myb 30*-related transcription factor. The present research work will provide rice breeders (i) the pre breeding material in the form of anticipated DSR adapted introgression lines possessing useful traits and alleles improving germination under deep sown DSR field conditions (iii) the base for the studies involving functional characterization and knocking of *Myb-30* gene. The next step is to understand the genetics and validate the molecular mechanism behind the cross-talk between ethylene-mediated signalling (*EIN-3*) and *Myb-30* in regulating mesocotyl elongation in rice using genome editing. We are at the stage of testing the cross-talk hypothesis through active research and development with analytical and laboratory studies.

## Materials and methods

The plant material of this study consisted of biparental mapping population constituting 420 recombinant inbred lines derived from crossing between PR126 (an early elite *indica* rice variety) and IRGC 128442 (an accession from 3K-RGP panel with better germination ability from deep sowing depth). The phenotypic characterization was carried out in the field and screenhouse areas of School of agricultural biotechnology, Punjab Agricultural University, Ludhiana, India. The six accessions selected from 3K-RGP including Aus344, N22, Kula Karuppan, NCS237, Ashmber and IRGC 128442 along with checks (PR121, PR126, PR128, PR129, PB1509) were evaluated first by drilling at 8 cm soil depth under direct seeded cultivation conditions under field conditions in 6 m plot in 3 replications with 0.2 m row to row and 0.2 m plant to plant spacing. The best donor was selected and used to develop the bi-parental mapping population in the background of PR126. The introgression of ‘desired seed vigour trait’ for early-uniform emergence from deeper soil depth under dry direct seeded cultivation conditions in background of PR126 was initiated in kharif 2019. A total of 200 F_1_s seeds were produced. The F_1_ seeds were advanced to F_2_ and F_1_ seeds were also backcrossed to PR126 in 2019-2020 rabi season under controlled conditions in screenhouse. The F_2_s were advanced to F_3_ under field conditions in 2020 Kharif season and the BC_1_F_1_s were backcrossed to PR126 to generate BC_2_F_1_s. The F_3:4_ and F_4:5_ populations were evaluated under screenhouse and field conditions in 2021 and 2022 *kharif* seasons, respectively. The BC_2_F_1_s were advanced to BC_2_F_2_s and then to BC_2_F_3_s in 2021 and 2022 *kharif* seasons, respectively.

### Phenotypic evaluation of mapping populations in screenhouse

A set of 750 breeding lines derived from the F_3:4_ population PR126/IRGC128442 was screened for the emergence from deeper soil depth under controlled screenhouse conditions. Based on 0-50 percent germination (212 lines), 50-80 percent germination (68 lines), and 80-100 percent germination (96 lines), a set of 420 breeding lines were selected. The F_3:4,_ and F_4:5_ and control checks including PR126, PR121, PR128, PR129 and PB1509 (low/no germination from deep sowing depth) Aus344, N22, Kula Karuppan, NCS237, Ashmber and IRGC 128442 (better germination from deep sowing depth) were screened for the emergence from deeper soil depth under controlled screenhouse conditions in 2021 *kharif* season and 2022 *kharif* season, respectively. The BC_2_F_2:3_ population constituting 339 plants along with above mentioned checks were screened for the emergence from deeper soil depth under controlled screenhouse conditions in 2022 *kharif* season.

### Screening for emergence from deeper soil depth in screenhouse

Six seeds of all the breeding lines was sown in the plastic trays filled with soil at 4 cm, and 10 cm depth (Figure 1). In each tray, the donor possessing the trait of interest (IRGC 128442) was kept as a positive check and PR126 was be kept as a negative check. The seed sown at 4 cm depth was act as a control for soil depth at 10 cm. The phenotypic data on percent germination at a different soil depth, uniformity of germination, days to germination, mesocotyl and coleoptile length, root and shoot length was recorded. The total number of emerged seedlings (percent germination) per breeding line was recorded daily starting from the 4 DAS (days after sowing) until the 15 DAS. The destructive sampling of all the six plants per breeding line was performed at 16 DAS to evaluate primary root and shoot traits. The total root length (RL) and shoot length (SL) was measured with a centimetre scale (cm) and mesocotyl/coleoptile length (ML) with vernier calliper for the six plants sampled per breeding line.

### Phenotypic evaluation of mapping populations under field conditions

The F_3:4_, F_4:5_ and BC_2_F_2:3_ populations were evaluated in the field under direct seeded cultivation conditions in an augmented design in 1.5 m paired row plot maintaining row to row spacing of 20 cm and plant to plant spacing of 20 cm in 2021 Kharif season and 2022 Kharif season, respectively. The observations on early vegetative vigour, days to 50% flowering, plant height, grain yield and related traits were recorded in both the years in F_3:4_, F_4:5_. The data on days to 50% flowering (DTF) was recorded in the field when 50% of the plants in the paired row plot exerted their panicles. The plant height (PHT) was measured as the mean height of three random plants, from the plant base to the tip of the highest panicle during the maturity stage. The panicle length (PL) was measured on centimeter (cm) scale and the number of panicles/plant (P/P) were counted manually. At physiological maturity, the harvested grains were threshed, dried and grain weight per breeding line was recorded in g (gram).

### Statistical analysis

The analysis of variance (ANOVA) and the year wise mean were calculated using mixed model analysis in PBTools V 1.4.0. The Fisher’s t-test was used to determine the significant difference among the breeding lines. The correlation analysis among different traits was performed in R. v.1.1.423.

### Genotyping and Variant calling

Genomic DNA of the F_3:4_ mapping population including PR126 and IRGC 128442 was prepared from 21 days old seedling. The integrity was analyzed on gel electrophoresis and then subjected to high throughput ddRAD sequencing using Illumina HiSEQ 4000. The dual enzyme (ddRAD) digested libraries was prepared and run on bioanalyzer for the library profiling. The raw sequences were generated. The paired-end sequencing and the read processing were carried out at NGB diagnostics Private Limited, New Delhi (India). The Illumina adaptor sequences were removed first and quality trimming of the adaptor-clipped reads was performed. The reads with final length of <20 bases were rejected from further analysis. The sequencing reads were mapped against the reference genome sequence of *O. sativa v7*.*0* (http://rice.plantbiology.msu.edu/pub/data/EukaryoticProjects/osativa/annotationdbs/pseudomolecules/version_7.0/all.dir/) using bwa (version 0.7.17-r1188). The read pairs with both read aligning in the expected orientation were used for further analyses. The SAMtools *version 0*.*1*.*1* (Li e*t al*., 2009) was used for the conversion of mapping files from the sam alignment format to the bam binary format. Picard software (*version 1*.*48*) was used to mark the duplicates marked in the sorted bam files. The generated bam files were used for the variant calling using unified Genotyper of Genome Analysis Toolkit, version 3.6 (GATK pipeline). The Vcftools (version 0.1.17) was used for the filtering of variant calls. Finally, the samples with MAF 5% and missing 10% were kept.

### Linkage map construction, QTL and fine mapping

The genotypic data of the mapping population with filtered SNP markers was used for the linkage map construction using JoinMap 4.1 software (Van Ooijen, 2006) and R/qtl package (Broman and Sen, 2009). The underlying algorithm used for calculating the QTL genotype probabilities and to deal with missing or partially missing genotypic data was hidden Markov model (HMM) (Baum et al., 1970) by EM algorithm (Dempster et al., 1977; Lander and Botstein, 1989). A permutation test using 1000 permutation (Churchill and Doerge, 1994) was used for determining the LOD threshold significance level. The QTL confidence interval (flanking markers) was determined based on 95% Bayesian Credible Interval method with an interval expanded to the nearest markers (Sen and Churchill, 2001; Manichaikul et al., 2006). Based on the highest LOD score in the LG, the peak marker for each QTL was identified using R/QTL’s ‘find.marker’ method. In addition, the mapping was also performed using QTL IciMapping software v4.1 (Meng et al., 2015). For the fine mapping, the SNPs between parents in the mapped genomic regions were identified from the already available whole genome resequencing data of the parents.

### Designing of KASP markers and KASP assay

A total 15 KASP (Kompetitive Allele-Specific PCR) markers in the identified genomic region on chromosome 7 associated with the traits improving germination of rice under deep sowing depth were designed following protocol as mentioned in Sandhu et al. 2022. The designed 15 KASP markers were used for the polymorphic survey of the donor (IRGC 128442) and the recipient backgrounds (PR126) used in the present study. The polymorphic markers were used to fine map the identified genomic region using BC_2_F_2:3_ population. The KASP genotyping assays were carried out following protocol as mentioned in Sandhu et al., (2022).

### Identification and validation of candidate genes

RNA isolation and sequencing: Mesocotyl and coleoptile sampling for both the varieties (PR126 and IRGC 128442) at three different sowing depths (4, 8 and 10 cm) was done at every alternate day, starting from 3 days after sowing (DAS) to 15 DAS. Mesocotyl and coleoptile length was measured using a vernier calliper and the data was recorded from 10 seedlings per treatment. The appropriate stage/timepoint when there were significant variations in mesocotyl and coleoptile elongation was observed, tissue samples for transcriptomics study were collected. The mesocotyl and coleoptile samples were collected with three biological replicates and immediately frozen in liquid nitrogen. Total RNA was isolated using TRIzol reagent (Chomczynski and Sacchi, 1987). Three technical replicates were prepared.

The 108 RNA samples {2 genotypes (PR126, IRGC 128442), 2 treatments (4, 10 cm) and 3 stages (5DAS, 10DAS and 15DAS), 3 biological replicates and 3 technical replicates) were used for RNA sequencing using the Illumina HiSeq 4000 platform at NGB Diagnostics Private Limited, New Delhi (India). The raw reads obtained were pre-processed by removing adaptor sequences and discarding empty reads and low-quality sequences. All clean reads obtained were assembled and aligned to the reference genome (MUS Rice Genome Annotation Project Release 7, http://rice.plantbiology.msu.edu) using BWA alignment tool followed by sorting.

### RNA-seq based Differential Expression analysis

The transcriptome based differential expression analysis was done using Transcriptomics Module of Omics box (version 2.0). A count table was generated using aligned sorted bam files. Transcripts was normalized using TMM (Trimmed mean of M values) and filtered based on CPM values less than 0.5. Pairwise Differential expression analysis was done for all the possible combinations using GLM (Likelihood ratio test) using edgeR software in omics box. Time course expression analysis was done to understand the behaviour of selected candidate genes using maSigPro software in omics box itself.

#### Candidate gene filtration and analysis

Transcripts having a false discovery rate (FDR) > 0.05 and log2 (fold change) < 2 was filtered and not considered for the further analysis. Functional analysis was performed on the selected candidate genes. DEGs (Differentially expressing genes) extracted from pairwise Differential expression analysis and time course expression analysis were blasted against the non-redundant protein sequence (nr v5) database using the cloud blast tool and used for GO (gene ontology) mapping and functional annotation against the Gene Ontology annotated proteins database, to obtain their functional labels through BLAST2GO software.

### Validation of the candidate genes

The shortlisted candidate genes were validated based on their expression profiles using a panel of 6 samples (PR126, PR121, a line with very short mesocotyl from the PR126/IRGC 128442 mapping population, IRGC 128442, NCS237, a line with long short mesocotyl from the PR126/IRGC 128442 mapping population) by performing RT-qPCR measurements.

## Author contributions

NS conceptualized this study, provided resources, compiled the results, and drafted the manuscript; APA, OPR, MP, MS, GP, and SG. conducted field and screenhouse experiments, APA and JS performed mapping studies, JS, OPR, SB, GA, and VKV conducted the RNA seq experiments, J.S. conducted the transcriptome analysis; TJ, GA, and AK conducted the RT-PCR studies; HP, MKS helped with RT-PCR studies; NS, RK, SK, and AK contributed to the critical revision of the manuscript.

## Additional information

### Supplementary Information

The all-supported data are available in supplementary material provided with the manuscript.

**Correspondence** and requests for materials should be addressed to N.S.

